# Hippocampal Astrocytes Encode Reward Location

**DOI:** 10.1101/2021.07.07.451434

**Authors:** Adi Doron, Alon Rubin, Aviya Benmelech-Chovav, Netai Benaim, Tom Carmi, Tirzah Kreisel, Yaniv Ziv, Inbal Goshen

## Abstract

Astrocytic calcium dynamics have been implicated in the encoding of sensory information, and modulating them has been shown to impact behavior. However, real-time calcium activity of astrocytes in the hippocampus of awake mice has never been investigated. We used 2-photon microscopy to chronically image CA1 astrocytes as mice ran in familiar or novel virtual environments and obtained water rewards. We found that astrocytes exhibit persistent ramping activity towards the reward location in a familiar environment, but not in a novel one. Using linear decoders, we could precisely predict the location of the mouse in a familiar environment from astrocyte activity alone. We could not do the same in the novel environment, suggesting astrocyte spatial activity is experience dependent. This is the first indication that astrocytes can encode location in spatial contexts, thereby extending their known computational capabilities, and their role in cognitive functions.

## Introduction

In recent years, groundbreaking research revealed many surprising roles for astrocytes in modulating neuronal activity, as well as behavior^1^. Intracellular astrocytic calcium elevations, a prominent signal in these cells, were vastly studied in-vitro and recent works had investigated them in-vivo as well (reviewed in ^2,3^). Different studies have shown that astrocytes respond to specific sensory stimuli with calcium transients in the visual cortex^4-7^, the somatosensory cortex^8-13^ and the olfactory bulb^14^. Anesthesia suppresses calcium signaling in astrocytes^15^, but only a minority of studies have investigated astrocyte activity in awake animals. Nevertheless, direct manipulation of astrocyte calcium signaling was shown to modulate behavior, thereby extending their role beyond sensory processing^e.g. 16-19^. Astrocytic calcium signals are also affected by the general state of the organism: They are elevated during arousal^7,8,20,21^, reduced during natural sleep^22,23^, and regulated by neuromodulators in-vivo^7,24,25^. However, the real-time calcium activity of astrocytes in the hippocampus of awake mice has not been investigated as of yet, let alone during performance of a multisensory cognitive task.

Place cells, a subset of pyramidal neurons in the hippocampal CA1 region, fire when the animal is in a specific location in space^26^ and are considered to be the neuronal underpinning of spatial memory. The neuronal representation of a given environment entails goal related information: When an animal navigates in a familiar environment, place cells exhibit over-representation of rewarded locations^27-30^ with narrower and more stable tuning curves than for other locations^31^. Furthermore, a subgroup of neurons was shown to represent the reward, independent of its location^32^. Upon exposure to a novel environment, the activity of CA1 place cells reconfigures to form a new map that is unique to that environment^33-37^, enabling neuronal discrimination between distinct contexts^38-40^. Recent works have also shown that subpopulations of inhibitory cells exhibit spatially tuned activity^41^, and are modulated by rewards^42^, but the role of astrocytes in this context has not been investigated.

Hippocampal astrocytes play an important part in memory processes, as shown by us and others^16,19,43^, hence we hypothesized that their activity will also be modulated during the performance of a spatial cognitive task. To explore the calcium activity of astrocytes in CA1 during a spatial paradigm, we used 2-photon calcium imaging of a population of astrocytes in this region as mice ran on a linear treadmill and navigated in a circular virtual environment to obtain water rewards. We show that astrocytes gradually increase their Ca^2+^ activity towards the previously learnt reward location when mice explore a familiar environment. Moreover, decoders using Ca^2+^ dynamics in populations of astrocytes as their input, accurately predicted the mouse location within the virtual environment. When mice were introduced to a novel virtual environment differing in visual and tactile cues, the astrocytic population was no longer modulated by reward location, suggesting that the activity elevation towards a rewarding location requires familiarity with the environment. Our results shed light on the computational capabilities of astrocytes, their role in contextual discrimination, and their contribution to cognitive functions.

## Results

### Two-Photon Calcium Imaging of Hippocampal Astrocytes During Navigation

Calcium imaging of astrocytes in subcortical brain regions was never performed in awake animals. To investigate the activity of a population of astrocytes as mice perform a spatial task, we combined 2-photon calcium imaging with a custom-made circular virtual reality apparatus, and trained head-fixed mice to run on a 170cm long linear treadmill belt to obtain water rewards. A circular virtual environment with multiple distinct visual cues was projected onto a curved screen in front of them (Figure 1A). Mouse locomotion on the treadmill was recorded and translated into movement along the virtual track. A single water reward was given upon completion of each 170cm lap, matched with a specific location in the virtual environment (Figure 1B).

**Figure 1:**
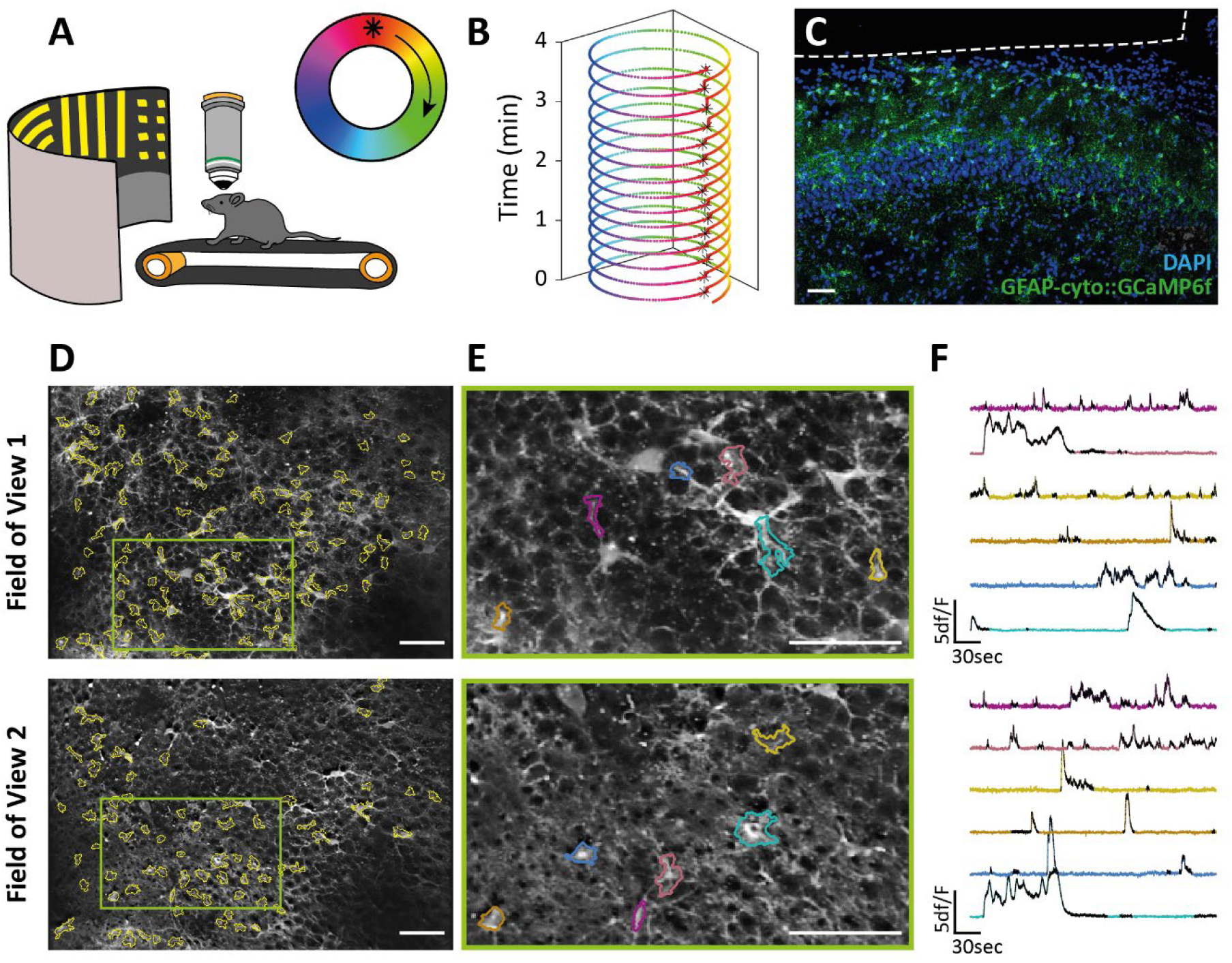
Ca^+2^ Imaging of CA1 Astrocytes in Mice Navigating a Virtual Reality. **A**. Experimental setup: Head-fixed mice ran voluntarily on a linear treadmill to proceed in a circular virtual environment projected onto a round screen in front of them to obtain water rewards, constantly given at a specific location (asterisk). **B**. Trajectory of a well-trained mouse, completing 15 laps in 4 minutes. The mouse typically stops only after a reward is given. Asterisks denote reward delivery. **C**. Expression of GCaMP6f (green) in astrocytes in CA1 following viral injection of AAV5-gfaABC1D-cyto-GCaMP6f. The imaging window was placed ∼100µm above the pyramidal cell layer, denoted by a white dashed line. **D**. Mean images of two fields of view acquired using a fast-z-tunable lens, showing numerous segmented astrocytic ROIs. **E**. Zoomed-in excerpts from the fields shown in D, with example ROIs and (**F**) their corresponding activity traces. Detected events are shown in black. Scale bars=50µm.

We virally expressed cytosolic GCaMP6f in dorsal CA1 astrocytes, (Figure 1C), enabling us to image real-time Ca^+2^ transients in astrocyte somata and main processes. To increase the number of astrocytic ROIs imaged in each session, we acquired separate time series from two fields of view (FOVs) using an electrical fast tunable lens focusing on different depths (Figure 1D-E, Movie 1). The obtained time series were motion corrected, and ROIs were semi-automatically segmented in each FOV separately. The signal from each ROI was extracted and the ΔF/F traces were calculated and used for binary event detection (Figure 1F).

### Astrocytes Increase Their Ca^2+^ dynamics Towards Reward Location

Reward locations are over-represented by place cells in familiar environments^27-31^, and a subgroup of neurons encodes rewards independent of their location^32^. However, it is unknown whether astrocytes exhibit location and reward specific responses. To tackle this question, we first examined the overall activity of the astrocytic population as mice traversed a familiar environment, and saw that it was characterized by synchronous activity epochs across many of the ROIs (Figure 2A, Movie 2). We calculated the number of concurrent events as a function of the mouse location, and saw that it increased towards the known reward location (Figure 2B). The number of concurrent events decreased during the stationary epochs that often occurred during reward consumption (Supplementary Figure 1A-B). The modulation of the astrocytic population activity by location was apparent across laps, and significantly different from shuffled data (correlation coefficient: 0.43±0.04, permutation test, p<0.05, n=4 mice) (Figure 2C-D, Supplementary Figure 1C-D; methods).

**Figure 2:**
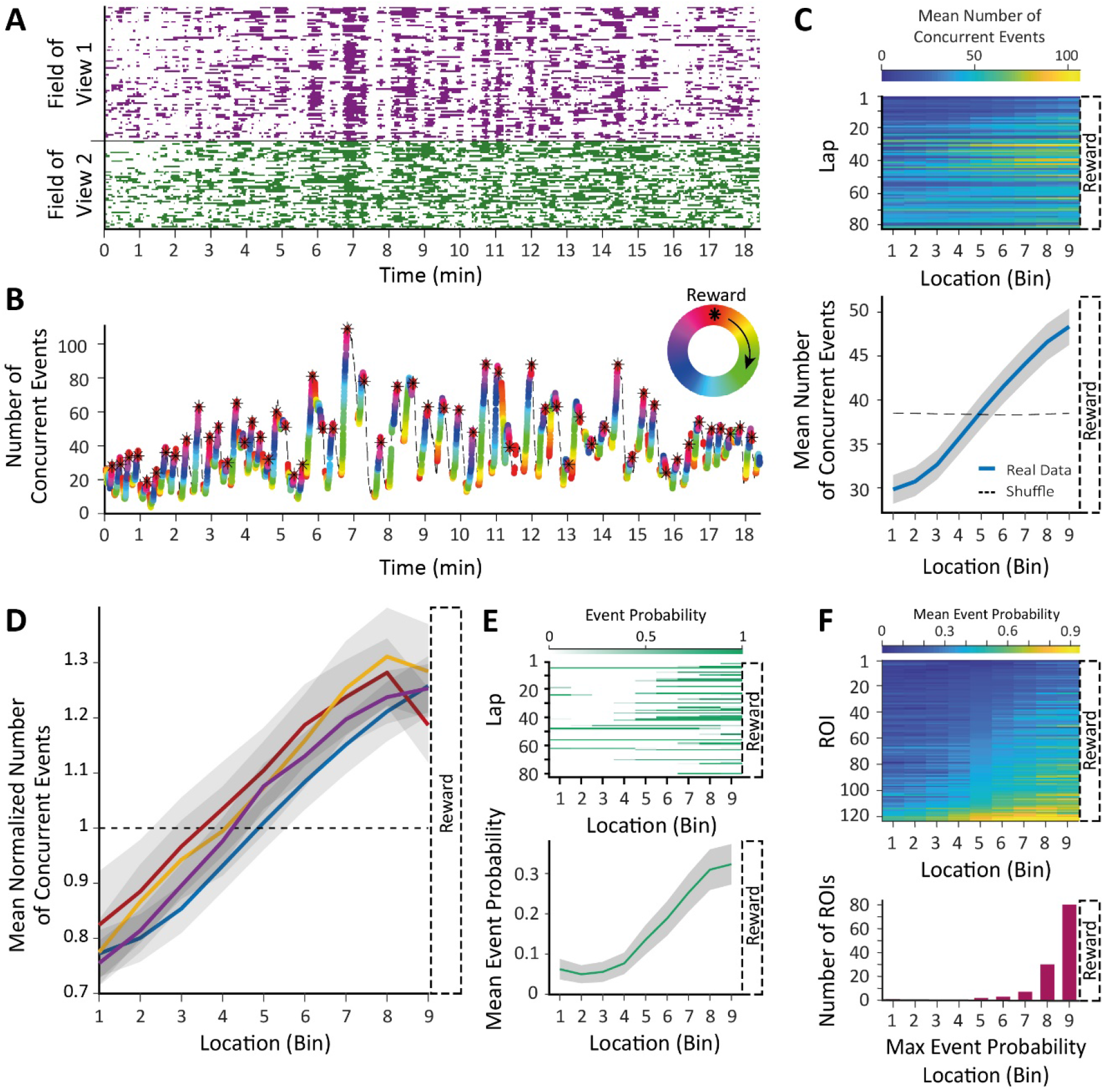
Astrocytic Activity Increases Towards Reward Location. **A**. Binary astrocytic Ca^+2^ traces acquired from ROIs in 2 fields of view when the mouse ran on the treadmill and advanced in the virtual reality. **B**. Summation of the binary traces shown in (A); The height is the number of concurrent events, and the colors denote mouse location along the track (the maze is depicted on the top right). The astrocytic activity ramps towards the known reward location, and decreases when the mouse is stationary. Asterisks denote reward delivery; dashed line denotes stationary epochs. **C**. The mean number of concurrent events of the mouse in (A-B) as a function of binned locations in all laps (top), and averaged across laps (bottom) is significantly different from shuffled data (correlation coefficient: 0.36, permutation test, p<0.01). **D**. Mean number of concurrent events as a function of location normalized by shuffled data in all 4 mice (blue is the one from C). The observed ramping was significantly different than the shuffled data (permutation test, p<0.05 in all 4 mice). **E**. Example of a ramping ROI, showing its event probability as a function of location and laps (top) and its mean event probability across laps (bottom). **F**. Mean event probability as a function of location of ROIs with significant spatial information obtained from 4 mice, sorted by the mean event probability in the central location bin (top). Most of these ROIs showed ramping, a gradual increase in mean activity probability apparent across laps, peaking near the known reward location (bottom). Data presented as mean (bold line) ±SEM (shaded area).

We next investigated whether single astrocytic ROIs have activation peaks covering the entire environment with over-representation of the reward location like place neurons. We found that about 30% (123/412, n=4) of the astrocytic ROIs had significant spatial information (methods). However, their activation peaks did not tile the entire track; the vast majority of these ROIs showed ramping, a gradual increase in mean activity probability apparent across laps, reaching its maximum near the known reward location (Figure 2E-F; methods), indicating that astrocytes represent the environment differently than neurons.

The increase in astrocytic Ca^+2^ events towards the reward may be the result of the mouse changing its velocity as it proceeds in the environment, as previous work has shown that astrocytes respond to locomotion in the cortex^21-23,44,45^ and the cerebellum^46^. To test this, we first looked at the number of concurrent events in the astrocytic ROI population as a function of velocity. The correlation between astrocytic activity and velocity was weaker than between location and astrocytic activity across laps (Supplementary Figure 1E-F). Moreover, when we investigated the interaction between location and velocity, we saw that the overall astrocytic activity varies more as a function of location in comparison to velocity (mean weighted STD: 0.07±0.01 and 0.03±0.01 for locations given velocity and velocities given location, respectively, paired t-test, t_(3)_=5.5, p<0.05)(Supplementary Figure 1G-H). Single astrocytic ROIs also exhibited higher variability across locations than across velocities (Supplementary Figure 1I). Taken together, these findings suggest that the astrocytic signal is modulated by location more than by the velocity of the mouse.

We show here for the first time that astrocytic activity in CA1 is modulated by location both when looking at single ROIs and at the entire imaged population. Unlike place neurons that fire selectively at specific locations throughout the entire space, astrocytes gradually increase their activity towards a single known rewarding location.

### Ramping of Astrocytic Activity Towards Rewarding Locations Requires Familiarity with The Environment

Previous studies have shown that CA1 place cells undergo global remapping upon exposure to a novel environment, and can discriminate between it and a familiar context^38-40^. Hippocampal astrocytes were never before imaged in awake behaving mice, and certainly not chronically, thus no such phenomenon is known in this population. Consequently, we asked whether ramping of astrocytic activity towards a rewarding location requires familiarity with the environment. To this end, we conducted chronic imaging of astrocytes both when mice navigated in a familiar environment, and when they were introduced to a novel one, differing in tactile and visual cues. We utilized fluorescent expression in sparse inhibitory neurons, allowing us to return to the same FOVs on subsequent days (Figure 3A). First, only repeated ROIs that were active on both sessions were included in the analysis (Figure 3B). As expected, the subpopulation of repeated astrocytic ROIs was significantly modulated by location in the familiar environment, gradually increasing its overall activity towards the reward location (Figure 3C). In the novel environment, however, this ramping was less apparent (correlation coefficient: 0.31±0.01 and 0.08±0.01 in the familiar or novel environment, respectively, permutation test for the correlation difference, p<0.05 in n=2 mice)(Figure 3D). Second, when all the imaged ROIs (not just the repeated ones) are included, the same result is found (correlation coefficient: 0.37±0.01 and 0.08±0.02 in the familiar or novel environment, respectively, permutation test for the correlation difference, p<0.05 in n=2 mice)(Supplementary Figure 2A-C).

**Figure 3:**
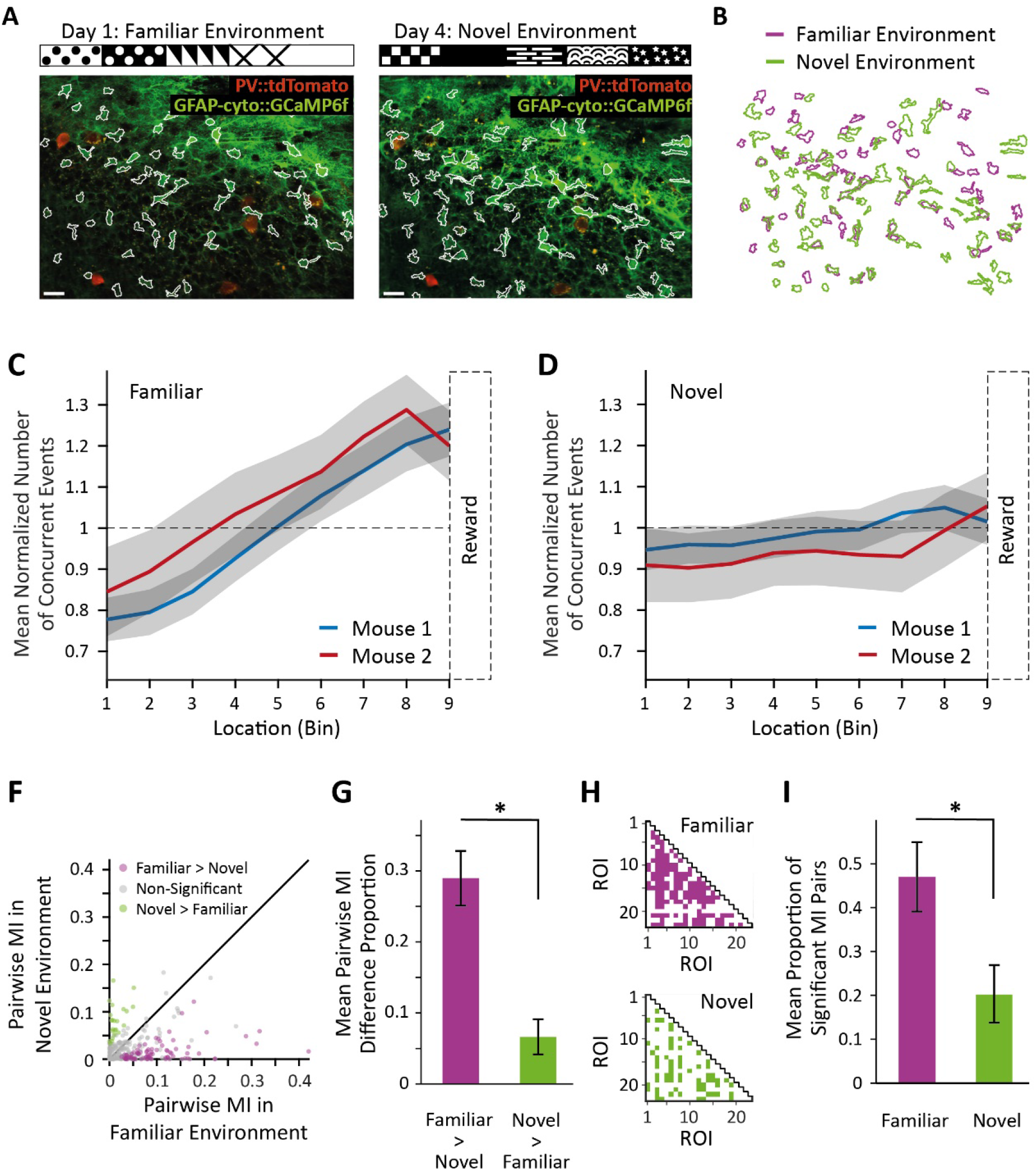
Astrocytic Activity Ramps Towards Rewarding Location in Familiar Environments, But Not in Novel Environments. **A**. Chronic imaging of the same field of view on consequent days when mice navigated in familiar (left) and novel (right) environments (Scale bars=25 µm; the walls of the virtual environments are drawn on top). The inhibitory neuronal tdTomato expression (in this case PV neurons) was used to ensure returning to the same field of view. ROIs were segmented independently on each session, and (**B)** their masks were registered to find repeated ROIs. Only ROIs that were active on both sessions were included in the analysis. The mean normalized number of concurrent events as a function of location in the familiar environment (**C**) and in the novel environment (**D**) for repeated active ROIs in 2 mice. The ramping of astrocytic activity is significantly larger in the familiar environment (permutation test, p<0.05, n=2). **F**. The pairwise MI between ROIs in the familiar and novel environments of the mouse shown in A-B. Significant differences are denoted in purple and green, for higher MI in the familiar or novel environment, respectively. **G**. A significantly larger proportion of ROI pairs had higher MI in the familiar environment compared to the novel environment (purple) than vice versa (green)(p<0.001). **H**. The ROI pairs that had significant MI in each environment of the mouse shown in A-B. **I**. The mean proportion of significant ROI pairs in the familiar environment is significantly higher than in the novel environment (p<0.001).

Next, we tested the mutual information (MI) between the astrocytic activity of ROI pairs that were repeatedly active in each context, and found pairs that had a significant difference between the familiar and novel environments (Figure 3F; methods). Significantly more ROIs decreased their MI in the novel environment compared to the familiar one (familiar>novel proportion: 0.29±0.04, familiar<novel proportion: 0.07±0.02, two proportions Z-test, p<0.001 in n=2 mice)(Figure 3G). Furthermore, the number of ROI pairs with significant MI in the familiar environment was higher than in the novel environment (proportion of significant MI ROI pairs: 0.47±0.08 and 0.2±0.07 in the familiar and novel environments respectively, two proportions Z-test, p<0.001 in n=2 mice)(Figure 3H-I; methods). Taken together, our results indicate that the astrocyte population discriminates between contexts, showing ramping towards the reward location and synchronous activity in familiar, but not in novel, environments.

### Mouse Location Can Be Decoded from Astrocytic Activity

We next asked whether the astrocytic population activity would suffice to determine the location of the mouse along the track. To this end, we constructed a linear-regression decoder for each mouse that predicted its location based on the binary Ca^+2^ activity traces, or on shuffled traces as control (Figure 4A-D, Supplementary Figure 3). The decoders estimated the mice trajectories significantly better than when tested on shuffled data (mean error size: 41.1±1.6cm for real data, 57.5±2.1cm for shuffled data; permutation tests, p<0.05 for all 3 mice)(Figure 4D, methods). Next, we asked whether similar linear decoders would be able to decode mouse location in the novel environment (Figure 4E-H). The performance of the decoders was not significantly different from when trained on shuffled data (mean error size: 51.2±4.7cm for real data, 55.5±0.9cm for shuffled data; permutation test, p>0.1 for both mice)(Figure 4H). Finally, we trained separate linear decoders to predict the normalized velocity of the mice from the binary Ca^+2^ activity traces, and found that their performance was not significantly different from when trained on shuffled data (mean error size: 0.2±0.02 for real traces, 0.2±0.02 for shuffled data; permutation tests, p>0.4 for all 3 mice)(Figure 4I-L), indicating that velocity cannot be decoded from the astrocytic activity. Taken together, our findings demonstrate that reconstruction of mice location trajectories from astrocytic activity can be done accurately using linear decoders, but requires familiarization with the environment, and, as opposed to neurons^36^, cannot be done in a novel environment.

**Figure 4:**
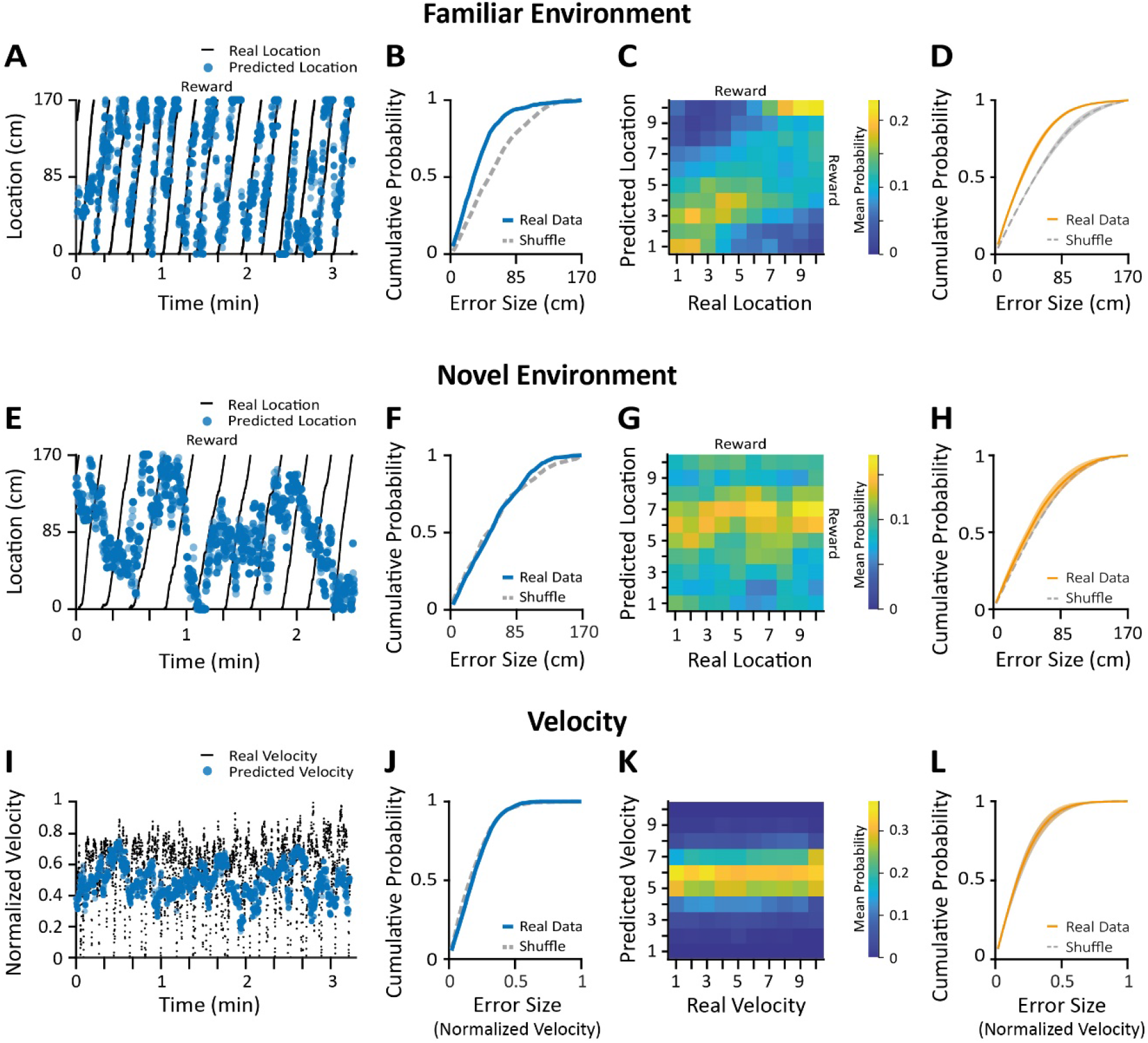
Accurate Decoding of Mouse Trajectory from Astrocytic Activity, Only in the Familiar Environment. **A-D**. A linear regression decoder was trained to predict mouse location in a familiar environment from binary astrocytic Ca^+2^ activity traces. **A**. An example trajectory of a mouse (black) overlaid with its accurately predicted location (blue dots). **B**. Error cumulative probability plot for the same data as in (A) showing the decoder trained on the real data is better than the one trained on shuffled data. **C**. Mean confusion matrix of the mouse shown in (A-B) based on 1000 linear decoder tests. The errors are concentrated on the diagonal, indicating that the predicted location is near the real location on average. **D**. Error cumulative probability of pooled data obtained from 3 mice, showing that the location decoders performed significantly better when trained on the real data than on shuffled data (permutation tests, p<0.05 for all 3 mice). **E-H**. A linear regression decoder was trained to predict mouse location in a novel environment from binary astrocytic Ca^+2^ activity traces. **E**. An example trajectory of a mouse (black) overlaid with its inaccurately predicted location (blue dots). **F**. Error cumulative probability plot for the same data as in (E) showing the decoder trained on the real data is not better than the one trained on shuffled data. **G**. Mean confusion matrix of the mouse shown in (I-J) based on 1000 linear decoder tests. The errors are concentrated on a horizontal line, indicating that the predicted location is independent of the real location on average. **H**. Error cumulative probability of pooled data obtained from 2 mice, showing that the location decoders did not perform significantly better when trained on the real data than on shuffled data (permutation tests, p>0.1 for all 2 mice). **I-L**. A separate decoder was trained to predict the normalized velocity of the mouse from the astrocytic Ca^+2^ activity traces. **I**. An example velocity trace of a mouse (black) overlaid with its inaccurately predicted velocity (clue dots). **J**. Error cumulative probability plot for the same data as in (I) showing the decoder trained on the real data is not better than the one trained on shuffled data. **K**. Mean confusion matrix of the mouse shown in (E-F) based on 1000 linear decoder tests. The errors are concentrated on a horizontal line, indicating that the predicted normalized velocity is independent of the real normalized velocity on average. **L**. Error cumulative probability of pooled data obtained from 3 mice showing the performance of the velocity decoders was not significantly better than when trained on shuffled data (permutation tests, p>0.4 for all 3 mice). Data presented as mean (bold line) ±SEM (shaded area).

## Discussion

This is the first report of astrocytic activity in the hippocampus of awake behaving animals to our knowledge. In this study, we chronically imaged CA1 astrocytes, as mice ran in familiar and novel virtual environments. While we notice no ‘place astrocytes’ that tile the whole environment, we find that astrocytic activity shows persistent ramping towards the reward location, but only when the context was previously learnt. Furthermore, we demonstrate that astrocytic population activity alone can be utilized to reconstruct mouse trajectories in familiar environments. To our knowledge, this is the first indication that astrocytes are involved in spatial tasks and can encode location in familiar contexts, thereby extending their known roles in cognitive functions.

Our data indicate that astrocytes encode position related information via gradual ramping towards the previously learnt reward location, both when examining the overall population and single ROIs activity. Such slow dynamics have been previously reported in neurons found in various brain regions^e.g.47-49^ that are involved in motor planning, working memory and decision making. Recently, a study has also shown that radial astrocytes of zebrafish gradually increase their Ca^2+^ activity as the animals learn the ineffectiveness of their actions, triggering a behavioral shift to passivity^50^. Our results show that calcium activity in CA1 astrocytes is elevated towards the reward location, consistent with a model in which astrocytic activity increases as evidence accumulates.

By chronically imaging astrocytes, we were able to compare their activity when mice navigated a familiar environment and when they were introduced into a novel one, differing in visual and tactile cues. Notably, in the novel environment the astrocytic activity was no longer modulated by location, and it did not suffice to accurately decode the mouse trajectory. Place cells rapidly emerge in CA1 following exposure to a novel environment^38,51,52^, and can be utilized to accurately decode the mouse location as early as the first lap^36^. Our results suggest that the astrocytic representation of an environment develops more slowly than the neuronal one. Importantly, astrocytic activity is significantly different between familiar and novel environments, which may indicate that astrocytes are involved in contextual discrimination, in conjunction with the neuronal representations^38-40^.

Astrocytes are known to have slow temporal dynamics (though see^12^), which may allow them to take part in computations that occur across long, behaviorally relevant time-scales. Moreover, the fact that they receive inputs from multiple neurons, may potentially allow them to serve as spatio-temporal integrators, as has been demonstrated in-vitro^53^ and in-vivo^50^. Astrocytes were previously shown to encode sensory stimuli with calcium transients in the cortex, and investigating the real time involvement of hippocampal astrocytes in various behaviors will deepen our understanding of cognitive functions and their underlying computations.

## Acknowledgements

We thank the entire Goshen lab for their support. AD is supported by the Azrieli fellowship and the ELSC graduate students scholarship. This project has received funding from the European Research Council (ERC) under the European Union’s Horizon 2020 research and innovation programme (grant agreement No 803589), the Israel Science Foundation (ISF grant No. 1815/18), and the Canada-Israel grants (CIHR-ISF, grant No. 2591/18). We thank Ami Citri, Yoram Burak, Yonatan Loewenstein, Eran Malach, Adi Kaduri Amichai, Adar Adamsky, and Adi Kol for the critical reading of the manuscript.

## Movie Legends

**Movie 1: 2-Photon Ca**^**+2**^ **Imaging of CA1 Astrocytes in Two Different Fields of View**. A video showing astrocytic Ca^+2^ activity in two fields of view, acquired using a fast-tunable lens focusing on different depths. Imaging in each field of view was acquired at 7.745 frames per second and is shown following motion correction. The video playback is sped up 4-fold the original acquisition rate.

**Movie 2: Ca**^**+2**^ **Imaging of Numerous CA1 Astrocytes During Virtual Navigation**. A video showing the Ca^+2^ dynamics in CA1 astrocytes (left panel) and the simultaneously recorded mouse location in the virtual environment (right panel). As the mouse approaches the known reward location, the astrocytic population gradually increases its activity. 126 ROIs were segmented in the complete dataset which includes an additional field of view (not shown), and their extracted signal is shown in figure 2A. Imaging in each field of view was acquired at 7.745 frames per second and is shown following motion correction. The video playback is sped up 4-fold the original acquisition rate.

## Supplementary Figures

**Supplementary Figure 1 (for Figure 2):**
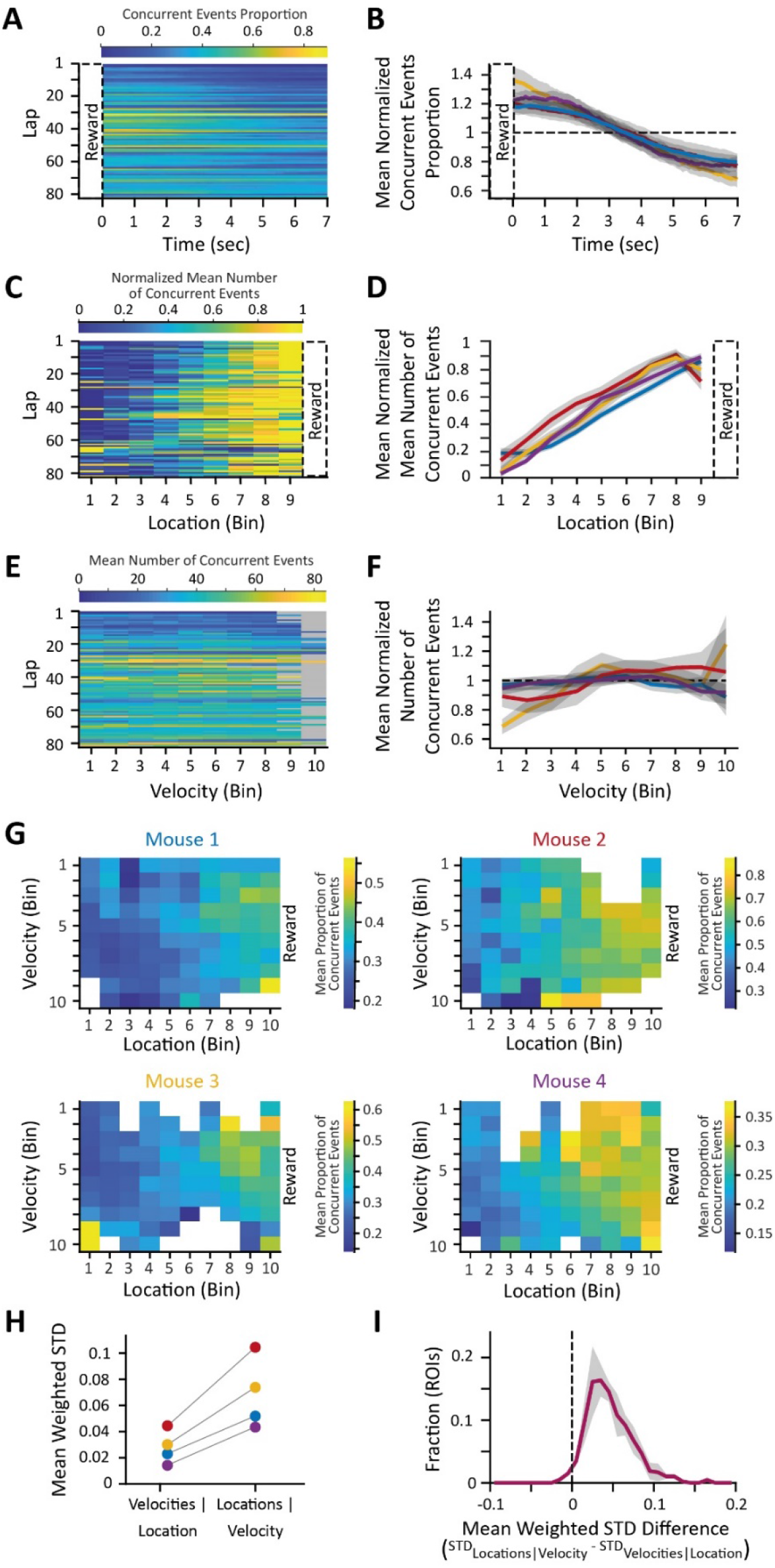
Astrocytic Activity Explains Location More Accurately Than Velocity. **A**. The proportion of concurrent astrocytic events following reward delivery in all laps of the same mouse shown in Figure 2A-C. **B**. Mean proportion of concurrent events as a function of time following reward delivery, normalized by mean concurrent events proportion in all 4 mice presented in Figure 2D (blue is the one from A), showing significant reduction over time (correlation coefficient: -0.46±0.07, permutation tests, p<0.01 in all 4 mice). **C**. The mean number of concurrent astrocytic events within each location bin, normalized within each lap, of the same data shown in Figure 2C. **D**. Mean normalized mean number of astrocytic concurrent events as a function of binned locations across laps in all 4 mice presented in Figure 2D (blue is the one from A), showing significant ramping (correlation coefficient: 0.74±0.04, permutation tests, p<0.01 for all 4 mice). **E**. The mean number of concurrent events of the mouse shown in Figure 2A-C as a function of binned normalized velocities in all laps. Gray bins denote no samples. **F**. Mean number of concurrent events as a function of binned normalized velocities, normalized by shuffled data in all 4 mice presented in Figure 2D (blue is the one from A), (correlation coefficient: 0.09±0.07, permutation test, p>0.05 in n=2 mice, p<0.05 in n=2 mice). **G**. The mean proportion of concurrent events as a function of location and normalized velocity in the 4 mice shown in Figure 2D. Ramping is apparent across locations, but not velocities. **H**. The mean weighted STD of the astrocytic population activity across locations for a given velocity (STD_locations|velocity_) is significantly larger than vice versa (STD_velocities|location_). **E**. The mean distribution of the difference between the mean weighted STD_locations|velocity_ and STD_velocities|location_ for single ROIs from the 4 mice shown in A-G. Most ROIs vary more across locations than across velocities. Data presented as mean (bold line) ±SEM (shaded area).

**Supplementary Figure 2 (for Figure 3):**
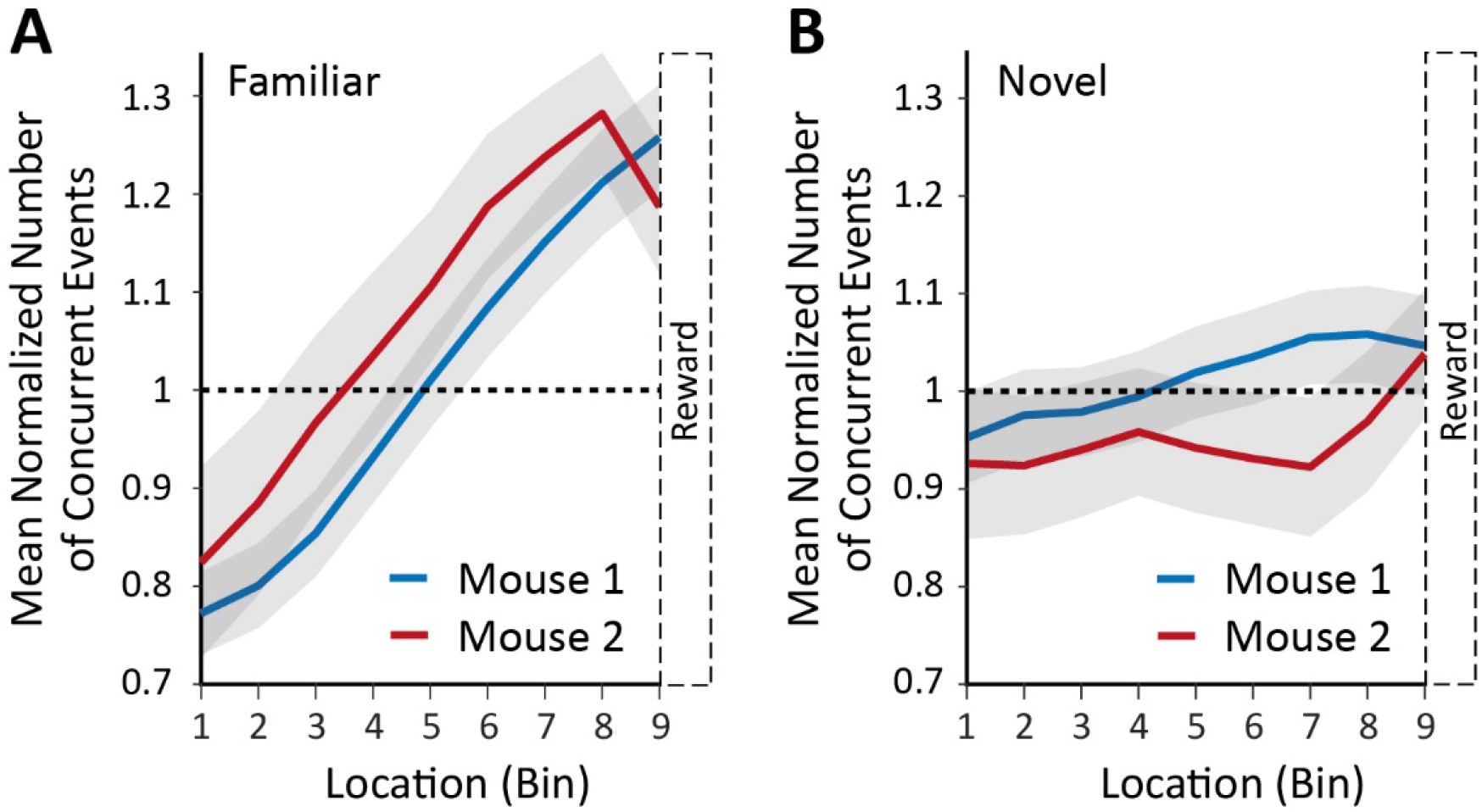
Astrocytic Activity Does Not Ramp Towards Rewarding Location in a Novel Environment. The mean normalized number of concurrent events as a function of location in the familiar environment (**A**, taken from Figure 2D) and in the novel environment (**B**) for all ROIs in 2 mice. The ramping of astrocytic activity is significantly larger in the familiar environment than in the novel environment (permutation test, p<0.05, n=2 mice). Data is for all ROIs, not just the repeating ones as in Figure 3.

**Supplementary Figure 3 (for Figure 4):**
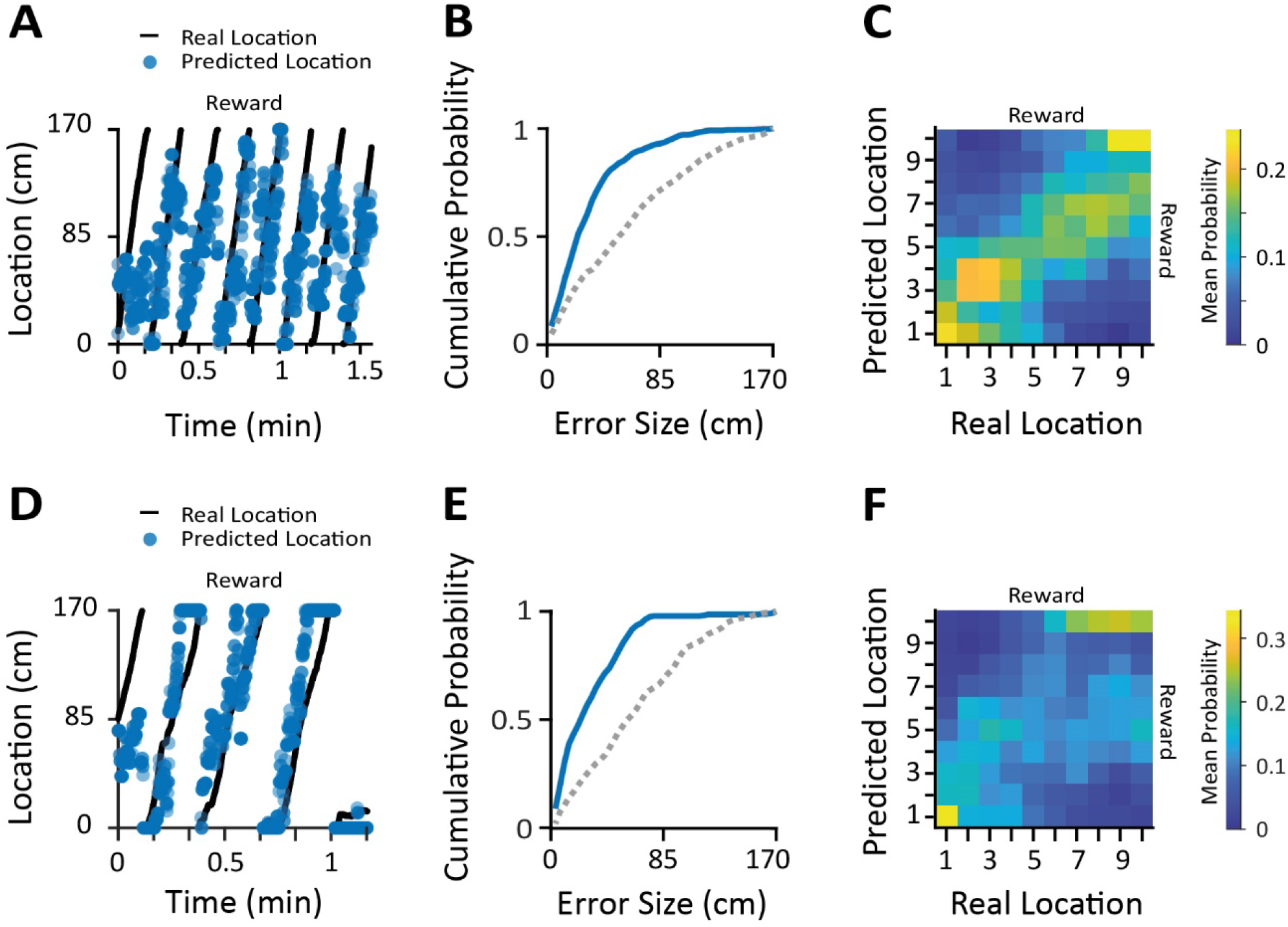
Accurate Reconstructions of Mice Trajectories from Astrocytic Activity in Familiar Environment. Additional trajectory reconstructions for 2 mice, that appear in the averaged data in Figure 4. **A**. An example trajectory of a mouse (black) overlaid with its accurately predicted location (blue dots). **B**. Error cumulative probability plot for the same data as in (A) showing the decoder trained on the real data is better than the one trained on shuffled data. **C**. Mean confusion matrix of the mouse shown in (A-B) based on 1000 linear decoder tests. The errors are concentrated on the diagonal, indicating that the predicted location is near the real location on average. (**D-F**) Similar to A-C, for another mouse.

## Supplementary Methods

### Animals

PV-tdTomato mice and an SST-tdTomato mouse were used for the experiments. The mice were generated by crossing *PV-IRES-Cre* (B6.129P2-Pvalb^tm1(cre)Arbr^/J, stock number 017320^54^) or SST*-IRES*-Cre (Sst^tm2.1(cre)Zjh^/J, stock number 013044^55^) mice with *Rosa-CAG-LSL-tdT (*Ai14; B6.129S6-Gt(ROSA)26Sor^tm14(CAG-tdTomato)Hze^/J, stock number 007908^56^) mice. 7-8 weeks old mice were housed on a 12 hr light/dark cycle in cages with running wheels. All mice were maintained under pathogen-free conditions in Tecniplast cages, on Teklad sani-chips (ENVIGO) bedding, at 20-24°C, and fed Teklad 2918SC (ENVIGO) pellets. Experimental protocols were approved by the Hebrew University Animal Care and Use Committee and met the guidelines of the National Institute of Health guide for the Care and Use of Laboratory Animals.

### Surgical Procedures

Mice were anesthetized with isoflurane, and their head placed in a stereotactic apparatus (Kopf Instruments, USA). The skull was exposed and a small craniotomy was performed. Mice were unilaterally microinjected 400nL viral vector using the following dorsal CA1 coordinates: Anteroposterior −1.85mm, mediolateral +1.4mm and dorsoventral −1.35mm from Bregma. All microinjections were carried out using a 10µl syringe and a 34 gauge metal needle (WPI, Sarasota, USA). The injection volume and flow rate (0.1μl/min) were controlled by an injection pump (WPI). Following each injection, the needle was left in place for 10 additional minutes to allow for diffusion of the viral vector away from the needle track, and was then slowly withdrawn. The craniotomy was sealed with bone-wax (Surgical Specialties, Tijuana, Mexico), and the exposed skull was covered with transparent super-bond (Sun Medical, Moriyama, Japan) for cementing an omega shaped head-bar (custom design, 3D printed) anteriorly to the craniotomy site. For postoperative care, mice were subcutaneously injected with Tramadex (5mg/kg).

Following at least one week of rest, mice were re-anesthetized with isoflurane in the stereotactic apparatus, and a biopsy punch (Kai Medical, Japan) was used to cut a ∼2.5mm diameter craniotomy over the injection site. Aspiration was used to remove the cortical tissue and top most fibers above the right dorsal CA1, and a glass cannula (2.4 mm diameter, 2.5 mm length, #0 cover slip bottom; self-fabricated) was inserted into the craniotomy. The skull was covered with opaque super-bond (Sun Medical, Moriyama, Japan) for cementing the cannula. An additional layer of dental acrylic was placed to minimize potential physical damage.

### Viral Vectors

pZac2.1 gfaABC1D-cyto-GCaMP6f (Addgene viral prep: #52925-AAV5; http://n2t.net/addgene: 52925; RRID: Addgene_52925).

### Immunohistochemistry

Mice were transcardially perfused with cold PBS followed by 4% paraformaldehyde (PFA) in PBS. The brains were extracted, post fixed overnight in 4% PFA at 4°C and cryoprotected in 30% sucrose in PBS. Brains were sectioned into 40μm thick slices using a sliding freezing microtome (Leica SM 2010R) and preserved in a cryoprotectant solution (25% glycerol and 30% ethylene glycol in PBS). Free-floating slices were washed in PBS, incubated for 1 h in blocking solution (1% BSA and 0.3% Triton X-100 in PBS) and incubated overnight at 4°C with primary antibodies (see below for a full list of antibodies) in blocking solution. Slices were then washed with PBS and incubated for 2 h at room temperature with secondary antibodies (see below for a full list of antibodies) in 1% BSA in PBS. Finally, sections were washed in PBS, incubated with 4,6-diamidino-2-phenylindole (DAPI; 1µg ml^−1^) and mounted on slides with mounting medium (Fluoromount-G, eBioscience).

#### Antibodies

The following primary antibody was used: Rabbit anti GFP (Novus, catalog no. NB-600-308; diluted 1:2000). Secondary Antibody: Donkey anti-rabbit conjugated to Alexa Fluor 488, (Jackson Laboratories, catalog no. 711-545-152; diluted 1:500).

### Confocal Microscopy

Confocal fluorescence images were acquired on an Olympus scanning laser microscope (Fluoview FV1000) using a ×10 air objective. Image analysis was performed using ImageJ (NIH).

### Linear Treadmill and Virtual Reality Apparatus

Fully awake mice were mounted on top of a linear treadmill with their head-bar secured to a custom-made holder under the microscope objective. The treadmill consisted of a 170cm belt with varying textures, circling 2 plastic wheels (custom design, 3D printed). To track mouse locomotion, rotations of a rotary encoder (S5-360-236-IE-S-B, US digital) placed in the frontal wheel were measured by Arduino boards. The locomotion data was synchronized with the microscope imaging frames, and translated into movement in the virtual environment. To compensate for sampling errors and belt stretches, an IR sensor connected to an Arduino board detected a white band on the inner side of the belt and auto-calibrated the VR accordingly on each lap.

A water solenoid valve connected to silicone tubes and a blunt 10cm needle delivered water rewards in response to TTL commands given by the VR computer via an Arduino board. Licking behavior was continuously monitored by a capacitance sensor (Atmel Microchip). To synchronize and digitize the valve, IR and lick signals with the imaging frames, we used a USB-6001 NIDAQ board (National Instruments) and acquired data at 500Hz using Matlab. The board recorded TTL signals from the microscope given on the beginning of each frame, as well as TTL commands sent to the valve and TTL inputs originating from the IR Arduino or lick detector on different analogue channels.

The virtual environments, designed using the Blender game engine, were projected onto a custom-made curved screen. Using a Java GUI, the specific environment of choice and reward locations were determined. The environments consisted of various visual patterns, in order to dissociate different locations in the virtual world. An Arduino board was used to trigger the initiation of the trial.

### Behavioral Paradigms

Mice were water-restricted and handled for 2-3 days, and then we began training them to run on the linear treadmill to obtain water rewards. Initially multiple water rewards were spread along the track, and as the mice improved on the course of 10-14 days, we gradually decreased the number of rewards until only one reward was present on each lap. We conducted the *familiar environment* using the same virtual environment and treadmill belt as in the training sessions. The *novel environment* experiment was conducted 3 days later using a different virtual environment and treadmill belt that the mice were not exposed to previously.

### Behavioral Analysis

We analyzed the behavior of the mouse using custom code run in MATLAB (mathworks). A lap was defined between each pair of rewards, and the relative location within it was calculated according to the rotary encoder tick count difference between the beginning and end of the lap. We smoothed the raw rotary encoder tick count using a moving average filter (∼0.3sec window) and defined movement epochs when the smoothed time series value was >1. Velocity was defined as the derivative of the smoothed time series.

### 2-Photon Microscope

2-Photon imaging was performed using the Neurolabware 2-photon laser scanning microscope (Los Angeles, CA, USA). Excitation light from a Ti:sapphire laser (Chameleon Vision II, Coherent, and then Chameleon Discovery TPC, Coherent) operated at 920 nm scanned the sample using a 6215 galvometer and a CRS8 resonant mirror (Cambridge Technology). Emitted fluorescence light was detected by GaAsP photo-multiplier tubes (Hamamatsu, H10770-40) after bandpass filtering (Semrock). XYZ motion control was obtained using motorized linear stages, enabled via an electronic rotary encoder (KnobbyII). We alternately scanned two imaging-planes with an electrically tunable lens for fast Z focusing (Optotune EL-10-30 NIR ETL; f=100 mm offset lens) to increase the number of astrocytic ROIs per session. A molding clay ring was mounted between the cannula and the objective in order to maintain the water reservoir and block external light. The Scanbox software, run on MATLAB, was used for microscope control and image acquisition. All images were acquired using a water immersion 16X objective (Nikon, 0.8 NA) with magnification of 2.8 or 3.4 to obtain 601×418µm or 516×366µm fields of view. The sampling rate was 15.49 frames per second, i.e. 7.745 frames per second for each imaged plane.

### Processing Ca^2+^ Imaging Data

#### Motion Correction

Ca^2+^ imaging movies were corrected for movement in each plane separately using either rigid motion-correction with the sbxaligntool algorithm (Scanbox) or non-rigid motion correction with the NoRMCorre algorithm^57^ in MATLAB (mathworks). When movements were still visible in specific frames, we removed them using the red channel. Specifically, we extracted the mean intensity time series of neuronal ROIs apparent in the red channel. Frames in which an ROI signal was >2 locally scaled MAD from the local median within a sliding ∼3.8sec window were excluded from the analysis.

#### ROI Detection

We used the sbxsegmenttool GUI (Scanbox) in MATLAB to semi-automatically detect ROIs based on the motion corrected movies. The segmentation was done separately for each field of view and for each session.

#### Signal Extraction

The mean fluorescence intensity time series were extracted from the segmented ROIs. The first 5 samples were removed from the analysis, as well as frames at the end of the movie if extensive bleaching was apparent. To synchronize the imaging data with the encoder data, we linearly interpolated the signal obtained from each plane. To obtain Δf/F time series for each ROI, we adapted previously published methods^58^ for the astrocytic signal. Specifically, we defined the baseline F as the 8^th^ percentile fluorescence value within a ∼125sec interval around each sample point. We then subtracted the baseline from each sample, and divided the result by the baseline F. Noisy ROIs, in which there were no apparent Ca^2+^ transients, were removed from the analysis.

#### Detection of Ca^2+^ Events

Potential events were first detected based on the Δf/F traces; For each ROI, we defined the event threshold as the sum of the mode fluorescent value and its distance from the mean minimal fluorescence value (based on the 100 smallest values). Next, we obtained potential events based on smoothed Δf/F traces (moving median, ∼3.9sec window), using the same calculation. We defined an event based on the smoothed traces, only if at least one of the samples within it was also detected as an event based on the original traces, and if it was >260ms long.

#### Registration of ROIs Across Sessions

To image the same FOV across sessions, we used the mean intensity images of each plane based on the motion corrected movies obtained during the first imaging session. We used the red channel in which inhibitory neurons were apparent to adjust the objective focus and optotune parameters until reaching the same FOV. The ROI masks obtained from each session were rigidly aligned using the sbxmatchfields function (Scanbox). Only ROIs that had >20% overlap, and were visually confirmed, were considered as repeated ROIs across days, and used for the comparison between familiar and novel environments unless otherwise stated.

### Astrocytic Activity Following Reward Delivery Analysis

We calculated the number of concurrent events as a function of time 7 seconds following reward delivery in each lap. To determine statistical significance, we permuted the signal using 1000 random cyclic shifts, and performed the same analysis as for the real data. To define significant modulation of astrocytic activity by location, we calculated the linear correlation between these variables from the real and shuffled data. Significance was determined when the correlation coefficient obtained from the real data was smaller than the 5^th^ percentile of the shuffled data correlation coefficient distribution.

### Modulation of Astrocytic Activity by Location Analysis

To obtain concurrent events and event probability maps, we discretized the location of the mouse along the track into 10 bins, each 17cm long. The last bin in which the reward was given and consumed was removed from further analysis unless otherwise stated. We also omitted frames in which the mouse velocity did not exceed the movement criteria (see previous section). We then calculated the mean number of concurrent events when examining the entire ROI population, or the event probability when examining single ROIs, per bin in each lap. To obtain the shuffled data activity maps, we permuted the signal using 1000 random cyclic shifts, and performed the same analysis as for the real data. The normalized number of concurrent events was calculated by dividing the real data mean number of concurrent events by the shuffled data pooled mean number of concurrent events. To define significant modulation of astrocytic activity by location, we calculated the linear correlation between these variables from the real and shuffled data. Significance was determined when the correlation coefficient obtained from the real data was larger than the 95^th^ percentile of the shuffled data correlation coefficient distribution.

### ROI Spatial Information Analysis

We calculated the spatial information of astrocytic ROIs using the event probability maps of each ROI, as previously described^59^:

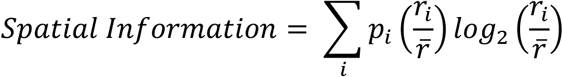

Where *r*_*i*_ is the Ca^2+^ event probability of the ROI given that the mouse is in the *i*^th^ bin; *p*_*i*_ is the probability of the mouse being in the *i*^th^ bin (samples spent in *i*^th^ bin/total session samples); 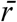 is the overall mean Ca^2+^ event probability; and *i* running over all the bins. We then performed 1000 cyclic permutations of the astrocytic signal, while keeping the behavior constant, and computed the spatial information for each shuffle. This yielded the p value of the measured spatial information relative to the shuffles.

### Mutual Information between Repeated ROIs Analysis

We calculated the mutual information between each pair of repeated ROIs in each environment, as well as the difference in mutual information between the 2 environments. We included all of the acquired astrocytic time series, during stationary and movement epochs of the mice. The pairwise mutual information was considered significant when it was higher than the 95th percentile value obtained from 100 cyclic permutations of the ROI signals. Mutual information difference was considered significant when it was larger than the 97.5th percentile or smaller than the 2.5th percentile obtained from 100 cyclic permutations of the ROI signals.

### Decoding Mouse Location and Velocity

To decode the mouse location or velocity based on astrocytic activity, we trained a linear regression decoder for each mouse separately. The signal was the binary astrocytic activity traces during movement epochs, and the output was the normalized location along the track or the relative velocity (i.e. between 0 and 1). We trained the decoder on 80% of the data, and tested its performance on the remainder. We ran 1000 train-test sets by dividing the data using a random cut-point. When the predicted location or velocity was out of the range [0, 1], it was trimmed (i.e. min(prediction, 1), max(prediction, 0)). To determine model significance, we created shuffled data by permuting the signal in a cyclic manner, and performed the same train-test procedure as for the real data. We computed the mean error (i.e. the mean difference between the real location and the predicted location) for the decoder trained on the real and shuffled data in each simulation. The model was defined as significant when <50 simulated datasets reached better performance than the real dataset (i.e. had a smaller mean error). Confusion matrices were calculated following binning of the real and predicted values into 10 bins.

### Statistical analysis

Data is presented as mean ± standard error of the mean (SEM) unless otherwise indicated. Sample number (n) indicates the number of ROIs or mice in each experiment and is specified in the figure legends. We performed permutation tests by conducting random cyclic shifts of the astrocytic data, and comparing the relevant distribution to the real data. We used Student’s t test to compare paired samples, and Z-test for binomial samples proportion comparison, as applicable. All the statistical details of experiments can be found in the result section. Statistical significance was considered at p<0.05. Analyses were performed using the IBM SPSS Statistics software (version 24) and Matlab.

## Notes

### Competing Interest Statement

The authors have declared no competing interest.

